# DNA supercoiling-induced shapes alter minicircle hydrodynamic properties

**DOI:** 10.1101/2023.01.04.522747

**Authors:** Radost Waszkiewicz, Maduni Ranasinghe, Jonathan M. Fogg, Daniel J. Catanese, Maria L. Ekiel-Jeżewska, Maciej Lisicki, Borries Demeler, Lynn Zechiedrich, Piotr Szymczak

## Abstract

DNA in cells is organized in negatively supercoiled loops. The resulting torsional and bending strain allows DNA to adopt a surprisingly wide variety of 3-D shapes. This interplay between negative supercoiling, looping, and shape influences how DNA is stored, replicated, transcribed, repaired, and likely every other aspect of DNA activity. To understand the consequences of negative supercoiling and curvature on the hydrodynamic properties of DNA, we submitted 336 bp and 672 bp DNA minicircles to analytical ultracentrifugation (AUC). We found that the diffusion coefficient, sedimentation coefficient, and the DNA hydrodynamic radius strongly depended on circularity, loop length, and degree of negative supercoiling. Because AUC cannot ascertain shape beyond degree of non-globularity, we applied linear elasticity theory to predict DNA shapes, and combined these with hydrodynamic calculations to interpret the AUC data, with reasonable agreement between theory and experiment. These complementary approaches, together with earlier electron cryotomography data, provide a framework for understanding and predicting the effects of supercoiling on the shape and hydrodynamic properties of DNA.

## INTRODUCTION

Nearly seventy years after Rosalind Franklin’s meticulous work that led to the first description of the structure of DNA (1), we are still working to understand how this remarkable molecule is organized (2), stored (3, 4, 5), activated (6, 7, 8, 9), and segregated into daughter cells. It is becoming increasingly apparent that negative supercoiling (the underwinding of the DNA double helix) provides a secondary or ‘hidden code’ that contributes to the three-dimensional (3-D) organization of the genome (10) that can be used by cells as a ‘molecular servomechanism’ to detect and regulate gene expression (11).

We recently discovered that the degree of curvature, dictated by DNA loop length, additionally tunes supercoiling-mediated effects and promotes mechanical crosstalk to expose DNA bases at specific distant sites (12). Exposed DNA bases drastically increase DNA flexibility to change the 3-D structure of DNA, which, conversely, influences the location and frequency of the disruptions to base pairing (12). Therefore, 3-D shape and base exposure are manifestations of supercoiling and looping (13). These findings underscore how supercoiling-dependent conformational changes may allow DNA to be an active participant in its transactions (13, 14).

We previously used minicircles of a few hundred base pairs and defined supercoiling to determine how supercoiling and looping modulates the 3-D structure of DNA (13). Although informative, these assays are laborious and only limited DNA sequences and buffer conditions have been explored (12, 13, 15). Electron cryotomography (cryoET) provides 3-D information on individual DNA minicircles of defined supercoiling (DNA topoisomers) (13), but the approach requires skill in the art, is time-consuming and the resulting structures are of low resolution. Increasingly powerful, atomic force microscopy can show the helical repeat of DNA as well as areas of base pair disruption (15), but the sequence is unidentifiable and the technique requires that DNA is adsorbed onto a flat surface. Not only might this adsorption distort DNA conformations but it means that they are visualized as 2-D projections with limited 3-D information.

Computational modeling would be of great value in helping predict DNA negative supercoiling and looping behavior, but thus far fails to account for supercoiling- and looping-mediated site-specific base exposure or resulting conformational changes. For example, most efforts at understanding looping (16) or cyclizing of DNA (17) ignore supercoiling. Therefore, new modeling efforts are needed, including the parameters of supercoiling and degree of curvature (dictated by DNA loop length). Before this modeling can be improved, however, detailed parameters of supercoiled loops of DNA must be determined.

Whereas cryoET is expensive and time-consuming, making it impractical to use for multiple conditions, AUC and electrophoresis rapidly assess properties in solution, and can be used to test multiple conditions simultaneously. Toward the goal of understanding how DNA sequence and 3-D shape are affected by negative supercoiling and looping, here we combined state-of-the-art AUC (18) with mathematical modeling to determine hydrodynamic parameters of supercoiled DNA minicircles. We derived partial specific volume (PSV) and anisotropy, and measured the sedimentation and diffusion coefficient for minicircle DNA of different degrees of negative supercoiling and lengths. With these values, we determined the density of DNA. We discovered that DNA length and supercoiling strongly affect the sedimentation properties of minicircle DNA but have no effect on the PSV.

We generalized the continuum elastic framework to accurately predict the previously observed DNA minicircle 3-D shapes (13). This generalization provides additional and complimentary information that will allow us to interpret supercoiling- and curvature-dependent DNA structural alterations. Emboldened by this accomplishment, we then combined the measured elastic and hydrodynamic properties of DNA minicircles using bead models and considering force and hydrodynamic effects to compute the hydrodynamic sedimentation and diffusion coefficients. These modeling results compared favorably to AUC measurements.

## MATERIALS AND METHODS

### Chemicals and reagents

MseI, Nb.BbvCI, Proteinase K, T4 DNA Ligase, low molecular weight DNA ladder, and 100 bp DNA ladder were purchased from New England Biolabs (Ipswich, MA, USA). Adenosine triphosphate (ATP), antifoam 204, dithiothreitol (DTT), ethidium bromide, and Rnase A were purchased from Sigma-Aldrich (St. Louis, MO, USA). Acrylamide, ampicillin, chloroform, and sodium chloride were purchased from Fisher Scientific (Pittsburgh, PA, USA). All other chemicals were purchased from VWR International (West Chester, PA, USA).

### Generation and purification of minicircle DNA

Plasmid pMC336 (13) was used to generate both the 336 bp and 672 bp minicircles via λ-integrase-mediated site-specific recombination as described (19). Double-length 672 bp minicircles contain two copies of the 336 bp minicircle sequence in tandem orientation and are a byproduct of the recombination used to generate 336 bp minicircle DNA.

### Generation of different DNA topologies

The “supercoiled” samples are the 336 bp or 672 bp minicircle products of the purification process. These were analyzed without further manipulation. To make nicked DNA, the minicircles were nicked at a single site using the nicking endonuclease Nb.BbvCI according to the manufacturer’s protocol. The 672 bp minicircle contains two copies of the BbvCI site and was thus nicked at both locations. Following nicking, the DNA was subsequently incubated at 80 °C for 20 minutes to inactivate Nb.BbvCI. Linear 336 bp was generated by incubating supercoiled 336 bp minicircle with MseI according to the manufacturer’s protocol. The linearized DNA was subsequently incubated at 65 °C for 20 minutes to inactivate the enzyme. “Relaxed” 336 bp minicircle DNA was generated by incubating the nicked minicircles with T4 DNA ligase in 50 mM Tris-Cl pH 7.5, 10 mM MgCl_2_, 1 mM ATP, and 10 mM DTT overnight at room temperature. “Hypernegatively supercoiled” 336 bp was generated in an identical manner as “relaxed”, except for the addition of ethidium bromide (6.5 *μ*g/ml) to the ligation reaction. Ligations were subsequently extracted with butanol (to both reduce the volume and to remove the ethidium bromide), extracted with chloroform, then precipitated with ethanol. The nicked, linear, and supercoiled minicircle samples were also subjected to butanol and chloroform extraction and ethanol precipitation in a similar manner to ensure that any differences observed could not be attributed to differences in how the samples were made. Following ethanol precipitation, DNA was resuspended in 50 mM Tris-Cl pH 8.0, 150 mM NaCl, and 10 mM CaCl_2_. DNA samples were subsequently subjected to multiple rounds of buffer exchange in the same buffer using a Amicon 0.5 ml centrifugal filter to ensure that buffer conditions were equal across all samples. DNA concentrations were determined using a Nanodrop spectrophotometer.

### Geometry and topology of DNA minicircles

DNA supercoiling is defined by the linking number (*Lk*), the total number of times the two single DNA strands coil about one another (20), thus it is an integer number by construction, if both strands are covalently closed. If one or both of the strands is not covalently closed, e.g., for nicked and linear DNA, *Lk* can adopt non-integer values. Another quantity determining the shape of DNA is the equilibrium helical repeat *h* defined as the number of base pairs between two locations where the backbones are aligned and is measured in base pairs per turn. The value of *h* is buffer dependent and ∼10.42 bp/turn in 10 mM CaCl_2_ (13). Using *h*, we can calculate the angle between terminal base pairs of a straight linear DNA segment of a given length *L*, which gives us the reference value *Lk*_0_ = *L/h*. Since *h* can, in principle, take any value, *Lk*_0_ is not restricted to integer values and usually has a fractional part. For relaxed 336 bp minicircles, we get *Lk*_0_ = 32.2, while for 672 bp *Lk*_0_ = 64.4. Therefore, the deviation from the most relaxed DNA structure is measured by the difference between *Lk*_0_ and *Lk* denoted by Δ*Lk* = *Lk*− *Lk*_0_, which for the relaxed configuration of the 336 bp minicircle yields Δ*Lk* = −0.2. Δ*Lk* is typically scaled to the DNA length to give the superhelical density *σ* =Δ*Lk/Lk*_0_.

Since *Lk* is constrained to integer values it is sufficient to report Δ*Lk* rounded to the nearest integer to uniquely identify experimental configurations, as used in Ref. (13). For the relaxed 336 bp minicircle, we round −0.2 to 0. For simplicity, we follow this convention when reporting experimental values in this work. At the same time, we keep track of the fractional parts to accurately compute the elastic properties of the minicircles.

### Gel electrophoresis

DNA samples were analyzed by electrophoresis through 5 % (for 336 bp and 672 bp samples) or 4 % (for 672 bp samples) polyacrylamide gels (acrylamide:bis-acrylamide = 29:1) in Tris-acetate buffer (pH 8.2) containing either 150 mM NaCl and 10 mM CaCl_2_ (5 % gels) or 10 mM CaCl_2_ (4 % gels) at 125 V (∼ 6 V/cm) for 8 hours. Buffer was continuously recirculated during electrophoresis. DNA samples were also analyzed by electrophoresis through 1.5 % and 3 % agarose gels (Seakem LE agarose, Lonza, Rockland, ME) in TAE (Tris-acetate + 1 mM EDTA) buffer at 100 V for 3 hours. Gels were subsequently stained with SYBR Gold (ThermoFisher Scientific, Waltham, MA), then visualized using a FOTO/ANALYST Investigator imaging system (Fotodyne, Hartland, WI, USA) with quantitation using ImageQuant TL, version 8.1 (GE Healthcare Life Sciences, Marlborough, MA, USA).

### Analytical ultracentrifugation

Linear, nicked, relaxed, supercoiled, and hypernegatively supercoiled 336 bp minicircles, and supercoiled and nicked 672 bp samples were measured by sedimentation velocity using an AN50Ti rotor in a Beckman Coulter Optima AUC at the Canadian Center for Hydrodynamics at the University of Lethbridge in Alberta, Canada. For details of minicircles used, see Supplementary Table 1.

All samples were measured in 50 mM Tris-Cl pH 8.0, 150 mM NaCl, and 10 mM CaCl_2_. 460 *μ*l of each sample at an absorbance (A) of 0.6 at 260 nm were loaded into cells fitted with sapphire windows and 12 mm double channel epon charcoal centerpieces (Beckman Coulter, Indianapolis, IN, USA). Data were collected in intensity mode at 260 nm, and at 20 °C at five different rotor speeds of 10, 14, 25, 35, and 45 krpm. After data collection at each speed was completed, AUC cells were thoroughly shaken to redistribute the minicircle DNA uniformly. Depending on minicircle topology and length, at 10 krpm, pelleting occurred between 559–770 scans, requiring 50–70 hours. At 45 krpm, pelleting occurred after 88–159 scans, requiring 2–4 hours. The density and viscosity of the buffer, estimated with UltraScan, was 1.00682 g/ml and 1.02667 cP, respectively.

### AUC data analysis

All data were analyzed with UltraScan-III, version 4.0 (6345) (18), using the UltraScan data acquisition module (21). UltraScan fits experimental data to finite element solutions of the Lamm equation, deriving distributions for sedimentation and diffusion coefficients (22, 23). Optimization is achieved by parallel distributed data analysis, which was performed on the UltraScan Science Gateway using XSEDE resources (Expanse, Bridges 2, Stampede), and high-performance computing clusters at the University of Montana and University of Lethbridge. The optimization process proceeds through a series of model refinement steps, which employs the two-dimensional spectrum analysis (2DSA) (24). This refinement process removes systematic noise contributions contained in the raw data and obtains exact boundary conditions (the radial positions at the meniscus and the bottom of the cells) as described in (25). The final 2DSA refinement result is used to initialize a genetic algorithm analysis (GA) (26), which is followed by a Monte Carlo GA analysis (27). The total concentration determined from each speed between identical samples was also compared to ensure no material was lost due to aggregation or degradation, and samples were comparable across all speeds for a global analysis. The Monte Carlo GA results from identical samples and different speeds were combined to initialize a global GA analysis over all speeds. UltraScan supports simultaneous fitting to datasets from multiple experiments performed at different speeds. A global analysis benefits from the enhanced signal of the diffusion coefficient at low speeds and the improved sedimentation signal at higher speeds (28, 29). This feature also enhances signal-to-noise ratios and improves the confidence limits for the determined hydrodynamic parameters. The global fitting algorithm in UltraScan is further explained in (18).

### Hydrodynamic properties

#### Translational diffusion coefficient

The translational diffusion coefficient *D* is inversely proportional to the translational frictional coefficient *f*,

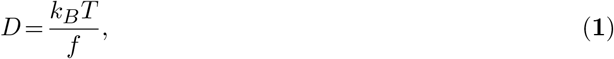

where *k*_*B*_ is the Boltzmann constant, and *T* the absolute temperature. In this work, diffusion occurs at very low concentrations of solute, which allows the analysis of transport coefficients in terms of single-particle properties only. For microscopic solid spheres of radius *R* suspended in a liquid of temperature *T* and viscosity *η* the Stokes-Einstein relationship reads

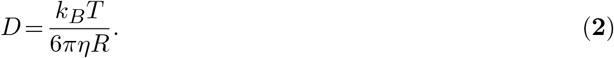

This relationship can be generalized to non-spherical molecules by introducing the effective hydrodynamic radius *R*_*h*_, defined as

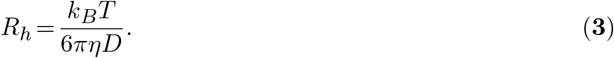

In aqueous solvents, macromolecules are typically hydrated, which adds to their apparent size and friction. The value of *R*_*h*_ derived from measured *D* includes these effects.

The hydrodynamic anisotropy of a given macromolecule was characterized by the frictional ratio *f/f*_0_, being the ratio of the measured frictional coefficient *f* and the frictional coefficient *f*_0_ of a spherical particle of the same volume. The anisotropy equals 1.0 for a spherical molecule and exceeds 1.0 for non-spherical molecular shapes.

#### Sedimentation coefficient

The sedimentation coefficient *s* depends on the molar mass *M*, the translational frictional coefficient *f* and the buoyancy of the particle, which is a function of its PSV, 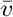, and solvent density *ρ*,

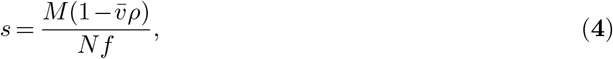

where *N* is Avogadro’s number. The Svedberg equation describes the ratio of the two parameters measured in a sedimentation velocity experiment, *s* and *D*, and provides a way to estimate the molar mass *M*, if the PSV is known:

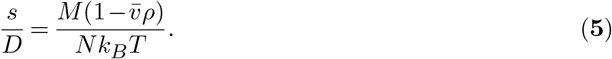

#### Apparent

*PSV* Eq. (**5**) considers a two-component system–an analyte with anhydrous molar mass *M* and a solvent with density *ρ*. However, our experimental solution also contains buffer components and ions that may be bound to the analytes. The degree of counterions bound to the analyte is dependent on solvent conditions and the ionic strength of the solvent, particularly for charged molecules (30). PSV is defined as the change in volume when one gram of analyte is added to the solvent, and is typically reported in units of ml/g. Because we do not know the precise amount of counterions bound to the analyte, we consider an apparent partial specific volume 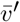, which can only be considered constant for a single solvent at a constant temperature and pressure. Rearranging the Svedberg equation allows the determination of the apparent PSV, provided the molar mass and the solvent density are known and the sedimentation and diffusion coefficients have been determined experimentally from a sedimentation velocity experiment

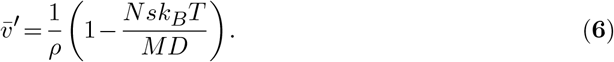

In our case, the molar masses are 207.576 kDa for the 336 bp minicircle and 415.152 kDa for the 672 bp minicircle, as calculated from the sequence (12) using molbiotools.com/dnacalculator. UltraScan automatically estimates the solvent density and viscosity from the buffer composition, adjusts the experimental *s*_*T,B*_ and *D*_*T,B*_ values to standard conditions (water at 20 °C) using the density and viscosity estimates from the buffer components, see (31, p. 117)

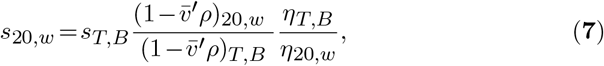

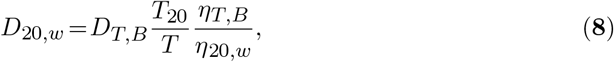

where *s*_*T,B*_ is the observed sedimentation coefficient at experimental conditions (temperature *T* = 293.15 K and buffer *B*). However, for the *s*_20,*w*_ corrections, the partial specific volume of DNA at standard conditions is required, but it is not known to us and impossible for us to measure. While literature values are reported for NaDNA (0.54 -0.55 ml/g) (32, 33), topoisomers here were studied in 10 mM calcium, which has a higher binding affinity to DNA than Na (34). Hence, we report here the experimentally measured values of *s* and *D* for all topoisomers, and the apparent partial specific volume under experimental conditions, 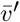, calculated by Eq. (**6**), and assuming a two component system.

### Finding equilibrium shapes of loops

To model the shapes of DNA minicircles with a given *Lk*, a variant of the Kirchhoff beam theory for inextensible rods (35) was used, which describes the twisting and bending of a uniform elastic filament of constant steric thickness *d*_*s*_, which was set to 20 Å in all computations. The helical repeat of the DNA yields a reference value of *Lk*_0_ = *L/h* for a given length *L*.

To model a DNA minicircle with a given *Lk* (and Δ*Lk*), we use an elastic beam representation in which a (closed) beam is characterized by two constants: bending rigidity *A* and geometric torsional stiffness *ω* (describing the cross-sectional shape, equal to 2/3 for circular cross sections). The energy density has quadratic contributions from the residual excess twist density Ω and local curvature *κ*. The total energy is thus given by

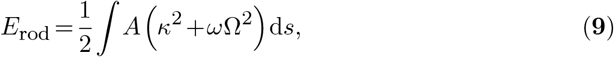

where Ω is computed from *Lk* and the shape of the filament centerline with the help of the Călugăreanu theorem (36)

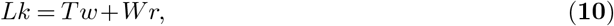

where twist (*Tw*) is defined as

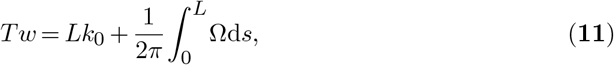

and writhe (*Wr*) is defined in the standard way (37). Scaling by *k*_*B*_*T*, the energy function can be made dimensionless, leaving the (width-to-length) aspect ratio *d*_*s*_*/L* and *ω* as the only parameters of the model. This approach was used by Coleman and Swigon (38) to categorize equilibrium shapes of looped filaments for a single aspect ratio *d*_*s*_ */L* = 8.2 ×10^−3^ (corresponding to a DNA minicircle of length 718 bp and *d*_*s*_ = 20 Å), which constitutes a benchmark for our computations. Coleman & Swigon began by solving the problem of a free beam segment subject to boundary conditions at each end, and two beams in contact along a contact line. Such solution fragments can be glued together at contact points to form a looped solution, subject to gluing conditions that ensure the continuity of the first two derivatives and appropriate jump conditions to account for beam-beam steric forces. This approach uses the same expression for the beam energy but addresses the energy minimization in a different way, either by solving an ordinary differential equation subject to appropriate boundary conditions when no contact forces are needed, or by direct minimization subject to no-overlap constraint when contact forces are present.

### Determination of critical Δ*Lk*

The stability of a computed minicircle shape depends on its Δ*Lk* (38, 39). For sufficiently small |Δ*Lk* |, a flat circular configuration is the only equilibrium solution. Upon increasing |Δ*Lk*|, at a thickness-dependent threshold value of critical (minimal) *Lk*_crit_, a figure-8 solution becomes admissible and the flat circular and figure-8 shapes coexist. Supercoiling further, above the thickness-independent threshold of 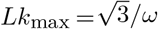, the flat circular shape is no longer a solution and only writhed configurations exist (39). For the prediction of *Lk*_crit_ for initial writhing of a minicircle, an approach based on solving an ordinary differential equation for centerline shape was used. The minimal value of |Δ*Lk*| required for writhing can be characterized by the existence of a configuration with a single contact point but with zero contact force. An ordinary differential equation was written for the beam centerline with minimum energy in a Cartesian parametrization subject to the boundary condition of a single contact point and no contact force and solved numerically using Mathematica (40), where boundary value problems are solved by the shooting method with conjugate gradient descent. For any given separation of the centerline at the self-contact location, one value of residual twist density was found. The relationship between the two was used to derive *Lk*_crit_(*d*_*s*_*/L*).

### Energy minimization of DNA minicircle shapes

Having determined the range of |Δ*Lk*| for which writhed configurations can be stable, the space of admissible configurations was examined and those configurations that minimized the elastic energy were investigated. Here, because of the presence of contact forces, a different numerical method was used. Representing the centerline shape with periodic cubic splines, direct energy minimization was performed over all possible shapes without self-intersections. Curves with 16 nodal points with enforced dihedral symmetry were subjected to a Monte Carlo minimization procedure. The bending energy was calculated directly from curvature using the adaptive Simpson’s algorithm and taking advantage of the twice continuously-differentiable nature of the cubic splines. The precise estimation of *Wr*, required to compute *Tw*, was performed by approximating the curve by 200 linear segments and using an algorithm proposed by Levitt (41) to deal with the singularities of the Gauss formulation. Steric interaction was introduced by tracking self-intersections through a large number (20*L/d*_*s*_) of sample points along the curve and a suitable steric energy penalty. Length constraint was imposed by computing the apparent length at each optimization step using the adaptive Simpson’s method and by imposing an energy penalty for the deviation from the prescribed length. Multiple sets of different control parameters for numerical optimization were tested to ensure both fast convergence and satisfactory precision. All final computations were done with identical discretization and penalty characteristics. Final values of penalty parameters, Monte Carlo procedure parameters and initial conditions are available on Github.

### Models for hydrodynamic radius

Solutions for Stokes flow around a slender toroidal object were developed by Johnson (42) that provide an asymptotic approximation in terms of slender-body theory. A fully analytical approach based on toroidal harmonics used by Goren & O’Neill (43) allows exact computations of all elements of the mobility matrix for a torus with an arbitrary aspect ratio. For a rigid axially symmetric particle of a given length *L* and hydrodynamic thickness *d*_*h*_, the mobility coefficients for translation along the symmetry axis and perpendicularly to it, *μ*_*z*_ (*L,d*_*h*_) and *μ*_*x*_(*L,d*_*h*_), respectively, are sufficient to compute the hydrodynamic radius, *R*_*h*_, by taking the inverse of their arithmetic mean

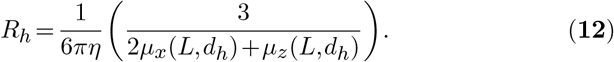

To theoretically determine *R*_*h*_ for an arbitrarily shaped molecule, a rigid bead model of its structure was constructed and its hydrodynamic radius was calculated using the ZENO software package (44, 45). In our case, the configuration of a minicircle was represented by 400 spherical and overlapping beads placed on the shape centerline, with diameters corresponding to the hydrodynamic thickness of the DNA molecule and the distance between overlapping beads summing up to the length of the molecule. This structure was then used to evaluate *R*_*h*_ for the composite particle. The diffusion coefficient at a given *T* and *η* is calculated from the definition of *R*_*h*_ in Eq. (**2**). The sedimentation coefficient is obtained from the Svedberg relation, Eq. (**5**).

## RESULTS

### Rationale

The study of DNA supercoiling and curvature has benefited from multiple complementary theoretical and experimental approaches. The combination of multiple approaches takes advantage of the knowledge gained from each approach while helping shore up their individual limitations.

We previously used 336 and 672 bp minicircles to study the effect of supercoiling and looping on DNA structure and have extensive 3-D structural data. AUC requires more material than cryoET (13), AFM (15), or other biochemical and biophysical analyses (12). We therefore tested supercoiled minicircle DNA as it was obtained from the bacterial cell instead of individual topoisomers, which are more difficult to isolate in large quantities. Additionally, a wide range of supercoiling was explored by testing relaxed, nicked, and hyper-negatively supercoiled samples. To determine the effect of circularity, linearized DNA samples were also analyzed.

We first characterized the minicircle samples by gel electrophoresis, which allows the topoisomer distribution of each sample to be precisely determined. Polyacrylamide gel electrophoresis effectively separates minicircle topoisomers and provides some insight into the conformational differences, although the theory underlying the differential migration is not fully understood.

We then applied advanced theoretical modeling to see whether it can explain the previously observed 3-D conformations of these minicircles (13). It was reasonably successful, and these advanced mathematical models could then be used to help analyze and interpret AUC data. Supercoiled DNA is a very structurally diverse molecule (13). Therefore, interpreting the AUC data using a simple model that approximates the shape as a sphere is not suitable.

### Electrophoretic characterization of DNA minicircles

DNA minicircles were analyzed by polyacrylamide gel electrophoresis. Both the helical repeat and conformation of DNA are sensitive to solution conditions (46, 47). Here we used 150 mM NaCl and 10 mM CaCl_2_, the same conditions used in analytical ultracentrifugation experiments.

Supercoiled topoisomers migrated much more rapidly on the polyacrylamide gel than relaxed topoisomers (Figure 1). In comparison, the different topologies had relatively similar mobilities on an agarose gel (Supplementary Figure 1). This difference in migration on polyacrylamide gels can be at least partially explained by the relative compactness of supercoiled minicircle conformations (13). The nicked and relaxed topoisomers had near-identical migration, suggesting that the single-strand break in the nicked minicircle does not significantly affect the global conformation. The lack of difference is explained by the number of helical turns in the 336 bp minicircle studied being close to a perfect integer value of 32 (under these conditions), resulting in the base pairs flanking the nick site being in close rotational alignment, allowing for favorable base stacking across the nick (12). We previously showed that when the rotational alignment is out of phase (i.e., when the number of helical turns deviates from a perfect integer value), the effect of a nick on polyacrylamide gel migration is much more pronounced (12).

**Figure 1.**
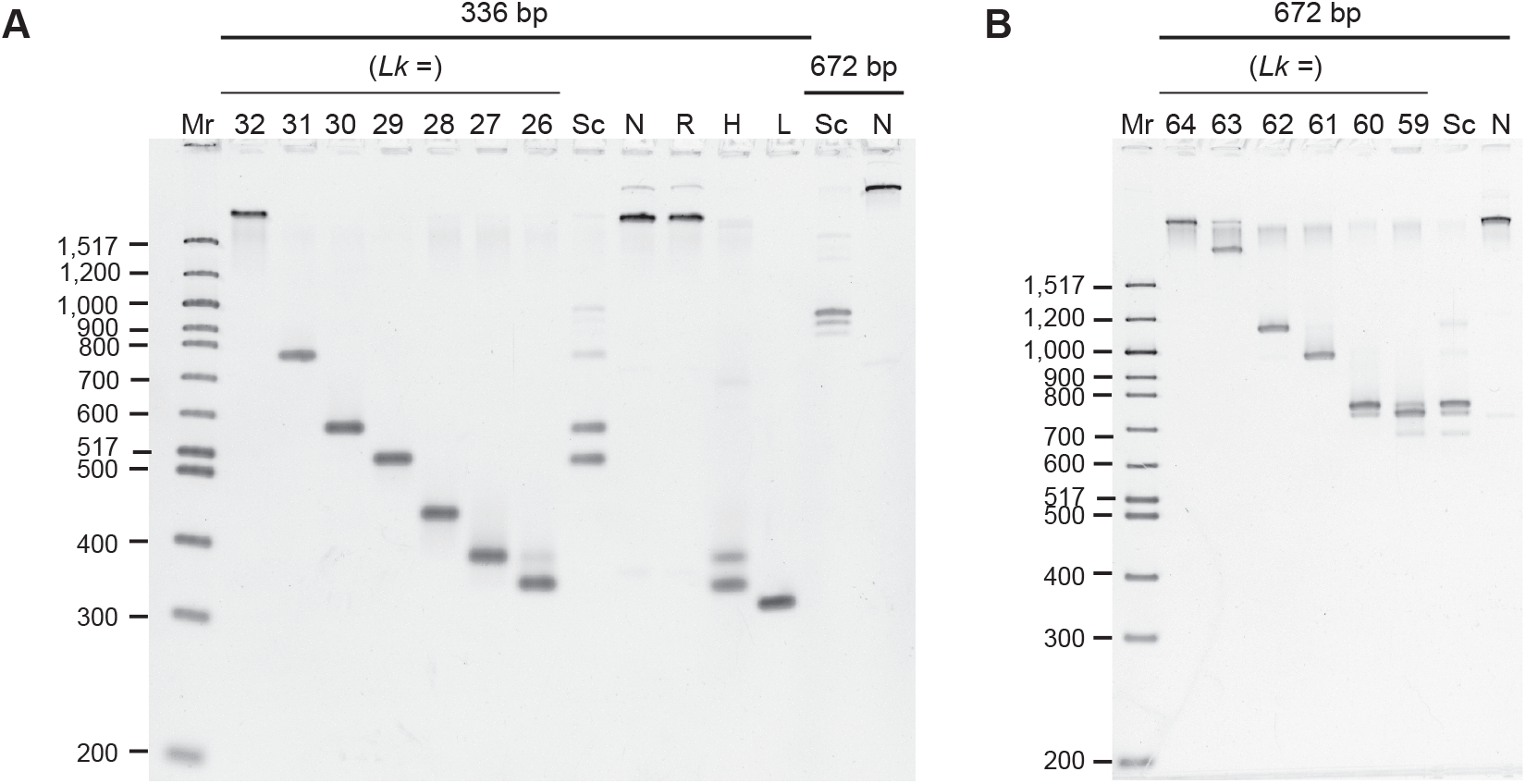
Electrophoretic mobility of minicircle DNA. **(A)** DNA samples were analyzed by polyacrylamide gel electrophoresis (5 % polyacrylamide) in 150 mM NaCl and 10 mM CaCl_2_ (the same conditions used in analytical ultracentrifugation). Mr: 100 bp DNA ladder, lanes 2–8: 336 bp minicircle topoisomer markers (*Lk* as indicated), lanes 9–13: 336 bp minicircle DNA samples (Sc: “supercoiled”, N: nicked, R: relaxed, H: “hypernegatively supercoiled,” L: linear), lanes 14–15: 672 bp DNA samples (Sc: “supercoiled,” N: nicked). **(B)** Determination of topoisomer identity in 672 bp samples. DNA samples were analyzed by electrophoresis on a 4 % polyacrylamide gel in the presence of 10 mM CaCl_2_. Mr: 100 bp DNA ladder, lanes 2–8: 672 bp minicircle topoisomer markers (*Lk* as indicated), lanes 9–10: 672 bp DNA samples (Sc: “supercoiled” as isolated from the bacteria, N: nicked).

The topoisomer distribution for the samples taken through to AUC analyzes was measured from quantification of digital images of the fluorescently stained gels using image analysis software. The “supercoiled” 336 bp sample contained primarily (48 %) Δ*Lk* = −3, 41 % Δ*Lk* = −2, and 7 % Δ*Lk* = −1 topoisomers. The sample also contained trace amounts of nicked 336 bp (1 %) and supercoiled 672 bp (3 %) minicircle DNA. The topoisomer distribution obtained (mean *σ* ∼−0.08) reflects the supercoiling level in the bacterial strain used to generate the supercoiling. The “hypernegatively supercoiled” sample contained primarily Δ*Lk* = −6 (61 %) and Δ*Lk* = −5 (33 %) topoisomers, with trace amounts of nicked 336 bp (4 %) and supercoiled 672 bp (3 %) minicircle DNA. This sample (with mean *σ* ∼ −0.15) is representative of the very high levels of dynamic supercoiling generated transiently during transcription and replication.

The supercoiled 672 bp sample contained primarily Δ*Lk* = −4 (63 %), Δ*Lk* = −5 (24 %), Δ*Lk* = −6 (6 %) topoisomers, and trace amounts of nicked 672 bp (2 %), Δ*Lk* = −2 (4 %) and Δ*Lk* = −3 (2 %) topoisomers. The identity of the topoisomers present in the supercoiled 672 bp sample was determined on a separate gel with 672 bp topoisomer markers. The topoisomer distributions of each sample are compiled in Supplementary Table 1.

### Analytical ultracentrifugation of DNA minicircles

Apparent PSVs obtained from a global multi-speed genetic algorithm-Monte Carlo analysis for each minicircle sample are summarised in Table 1. The derived PSV values did not show any apparent pattern that would indicate a dependence of the PSV on topoisomer conformation, and resulted in a near constant value of 0.482*±* 0.011 ml/g over all tested minicircles (see Table 1).

**Table 1.** Apparent partial specific volume for DNA minicircle topoisomers in the buffer. ^a^Determined by global sedimentation velocity analysis using the known molar masses. ^b^For all the 336 bp and 672 bp minicircle DNA species. ND, not determined.

| Partial specific volume<sup>a</sup> in mg/l |  |  |
| --- | --- | --- |
| Sample | 336 bp | 672 bp |
| Linear | 0.479 | ND |
| Relaxed | 0.470 | ND |
| Nicked | 0.469 | 0.495 |
| Supercoiled | 0.488 | 0.494 |
| Hypernegatively supercoiled | 0.479 | ND |
| Average value<sup>b</sup> | 0.482 ± 0.011 |  |

Using the determined average PSV, a frictional ratio *f/f*_0_ was derived from the sedimentation and diffusion coefficients obtained in the global analysis. Plots of the frictional ratio as a function of sedimentation coefficient, and the diffusion coefficient as a function of the sedimentation coefficient are shown in Figure 2. DNA topology had a significant effect on sedimentation and diffusion coefficients. In contrast to polyacrylamide gel electrophoresis, for which linear migrates fastest (Figure 1), circular molecules (nicked, relaxed, supercoiled and hypernegatively supercoiled) all sedimented faster than linear. AUC was additionally able to differentiate relaxed, supercoiled, and hypernegatively supercoiled samples. The supercoiled or hypernegatively supercoiled samples containing a mixture of topoisomers behaved as single species in AUC.

**Figure 2.**
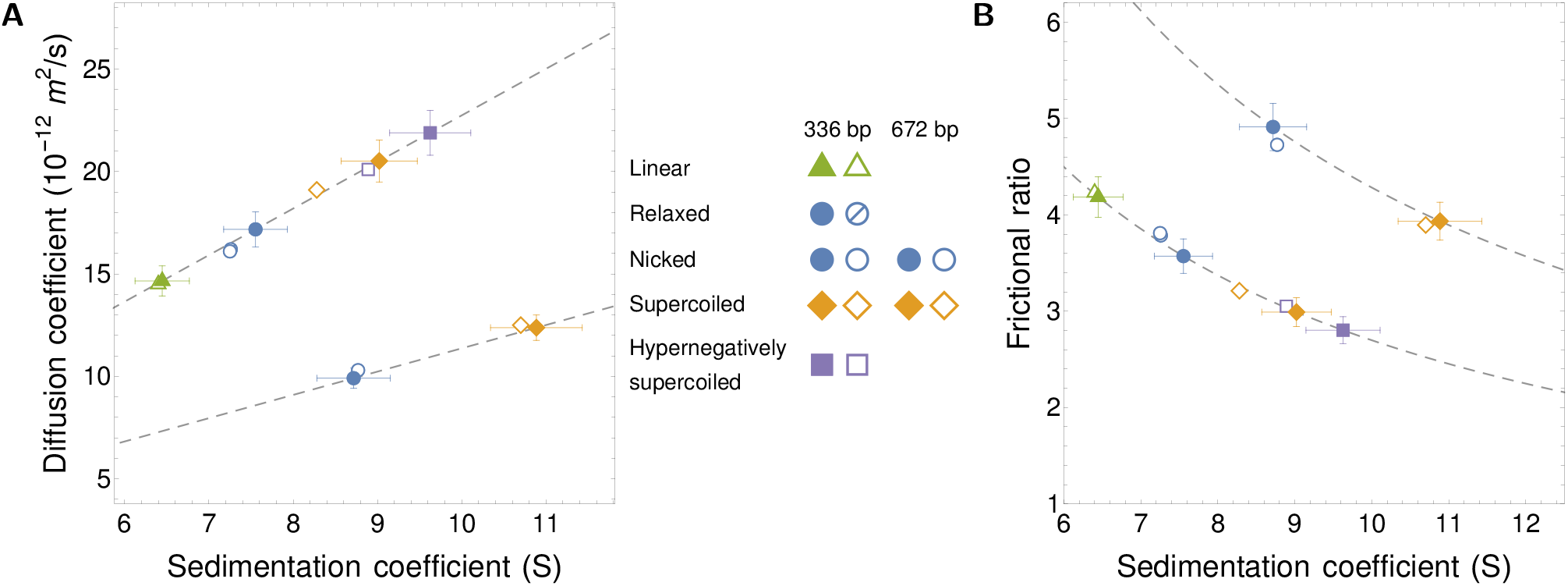
Measured and predicted diffusion and sedimentation coefficients for DNA minicircles. AUC measurements using global Monte Carlo-Genetic Algorithm analysis are marked as empty symbols. Theoretical predictions are presented with filled symbols. **(A)** Diffusion coefficient as a function of the sedimentation coefficient for topoisomers of 336 bp (upper branch) and 672 bp (lower branch) minicircle DNA. Dashed lines represent the constant mean value of PSV determined from AUC experiments on which theoretical predictions of the sedimentation coefficient are based. Experimental and theoretical data for 336 bp and 672 bp relaxed and nicked minicircles overlay almost completely and thus are impossible to discern in the plots. **(B)** Frictional ratio as a function of the sedimentation coefficient. Sedimentation coefficients *s* are measured in svedberg units (S), with 1 S = 10^−13^ s.

### Predicted shapes of DNA minicircles

Before employing our mathematical models to predict the effect of supercoiling and curvature on the hydrodynamic properties of DNA, we needed to first test how well these models predicted known equilibrium 3-D minicircle shapes. We adopted the strategy of building a coarse-grained representation of the equilibrium shapes of the minicircles obtained using our energy minimization codes. These models are reduced representations of macromolecules still capable of retaining key physical aspects (48). This approach has been highly successful in calculations of biomolecule properties in solution (49). Having found the shapes of DNA minicircles, we calculated their hydrodynamic radius for each conformation. The hydrodynamic radius was used to calculate the sedimentation and diffusion coefficients. We also tested our models against the 3-D structures of these minicircles, previously observed experimentally (13). Our aim was to develop a practical predictive theoretical framework to determine the measured transport coefficients.

#### Electrostatic screening and Brownian contributions

DNA molecules have substantial charge, which can be exploited, e.g., in electrophoretic measurements of DNA of different lengths. A qualitative comparison of the elastic and electrostatic forces is possible by considering scaling arguments. The Debye-Hückel equation is a well-established model of electrostatic interaction in a buffer containing counterions (50). In this approach, the interaction potential decays exponentially with separation due to screening. The decay rate is quantified by a characteristic distance, the Debye length (*R*_*D*_). Comparing *R*_*D*_ with the typical distances between base pairs gives a crude estimation of the influence of electrostatic forces. We estimated *R*_*D*_ for our setup to be 1.45 Å from the ionic strength of 230 mM using an ionic strength-based estimate 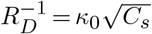, where *C*_*s*_ is the molar salt concentration in moles per litre and *κ*_0_ = 0.329 Å^−1^l^1*/*2^mol^−1*/*2^, as reported by Lim et al. (51). Thus the Debye length is much smaller than an average distance between different segments of the DNA molecule. Notably, this value of *R*_*D*_ was also much lower than that of earlier work, such as 30 Å in Ref. (52) for different buffer conditions.

The persistence length *P* of polyelectrolytes is the sum of two contributions,

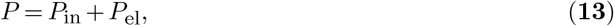

where *P*_in_ is an intrinsic persistence length due to the rigidity of the backbone, and *P*_el_ is an electrostatic persistence length, which accounts for buffer-dependent repulsion between neighboring ionic sites (53). The latter can be related to the Debye length as *P*_el_ = (4*κ*_2_*l*_*B*_)^−1^, with *l*_*B*_ being the Bjerrum length, according to the Odijk-Skolnick-Fixman theory (54, 55, 56). For linear or relaxed double-stranded DNA at room temperature, the persistence length in 0.1 M NaCl is approximately 500 Å (150 bp) (57). Based on persistence length, one can define effective bending and twisting energies for a circular shape as

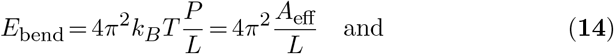

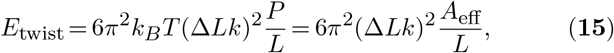

which, via Eq. (**13**), include both elastic and electrostatic contributions. Here, *A*_eff_ = *k*_*B*_*TP* is the effective bending rigidity of the DNA. The remaining long-ranged electrostatic contribution *E*_lr_ can be estimated as the interaction energy between *N* equal charges *q* at a typical distance comparable to the radius of the loop, *L/*2*π*, which amounts to

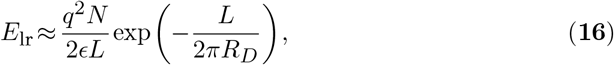

where *c* is the dielectric permittivity of the buffer. Note that these estimates do not take into account short-range electrostatic interactions between distant parts of the DNA that may come close together when supercoiled. For a circular loop with |Δ*Lk*| = 2 and equivalent length of 336 bp, we found *E*_bend_ ≈17*k*_*B*_*T, E*_twist_ ≈105*k*_*B*_*T* and a negligible value of *E*_lr_. The effective bending and twisting energies considerably exceed the typical energy of thermal fluctuations, *k*_*B*_*T*, which is somewhat surprising given that the minicircles are longer than *P*. This result may be a consequence of the relatively high energy stored in the form of bending energy *E*_bend_, which renders the circular configuration less prone to Brownian shape disturbances than a torsional-stress-free linear DNA. We therefore use the stiff beam approximation to describe minicircle shapes.

#### Critical ΔLk for writhing

The shape of a DNA minicircle depends primarily on |Δ*Lk*| and the width to length ratio *d*_*s*_*/L*. To predict 336 bp and 672 bp minicircle conformations for Δ*Lk* = 0 to |Δ*Lk*| = 5, we developed a theoretical framework that assumed that DNA can be modelled by a continuous elastic beam of Δ*Lk*, with the steric thickness *d*_*s*_ and stiffness *A* as determined by the inter-phosphate distance and persistence length, respectively. The previous theoretical study of Coleman & Swigon (38) focused on a circularised 718 bp DNA fragment and determined the structure of possible stable configurations, describing the contact diagrams in great detail. Our results were consistent with their findings for a particular thickness to length ratio (*d*_*s*_*/L* = 0.0082) but were applied to an arbitrary aspect ratio.

In Figure 3A we present the shapes obtained by our minimization procedure. Close to Δ*Lk* = 0, circular configurations were stable. At a critical value of |Δ*Lk*|, a figure-8 shape became energetically favorable, and the loop writhed to relax excess twist. For higher |Δ*Lk*|, the number of self-contacts increased, leading to a more writhed configuration. In this process, regions of high and low curvature emerged along the loop, as sketched in Figure 3B, with higher curvature present for higher |Δ*Lk*| s, as expected intuitively. For open configurations, the curvature was nearly constant and equal to 2*π*. The position along the loop changed from 0 to 1 and was measured from the point of contact or (in the case of multiple contacts) from the center of symmetry. With increasing negative Δ*Lk*, regions of increased curvature appear far away from self-contact points.

**Figure 3.**
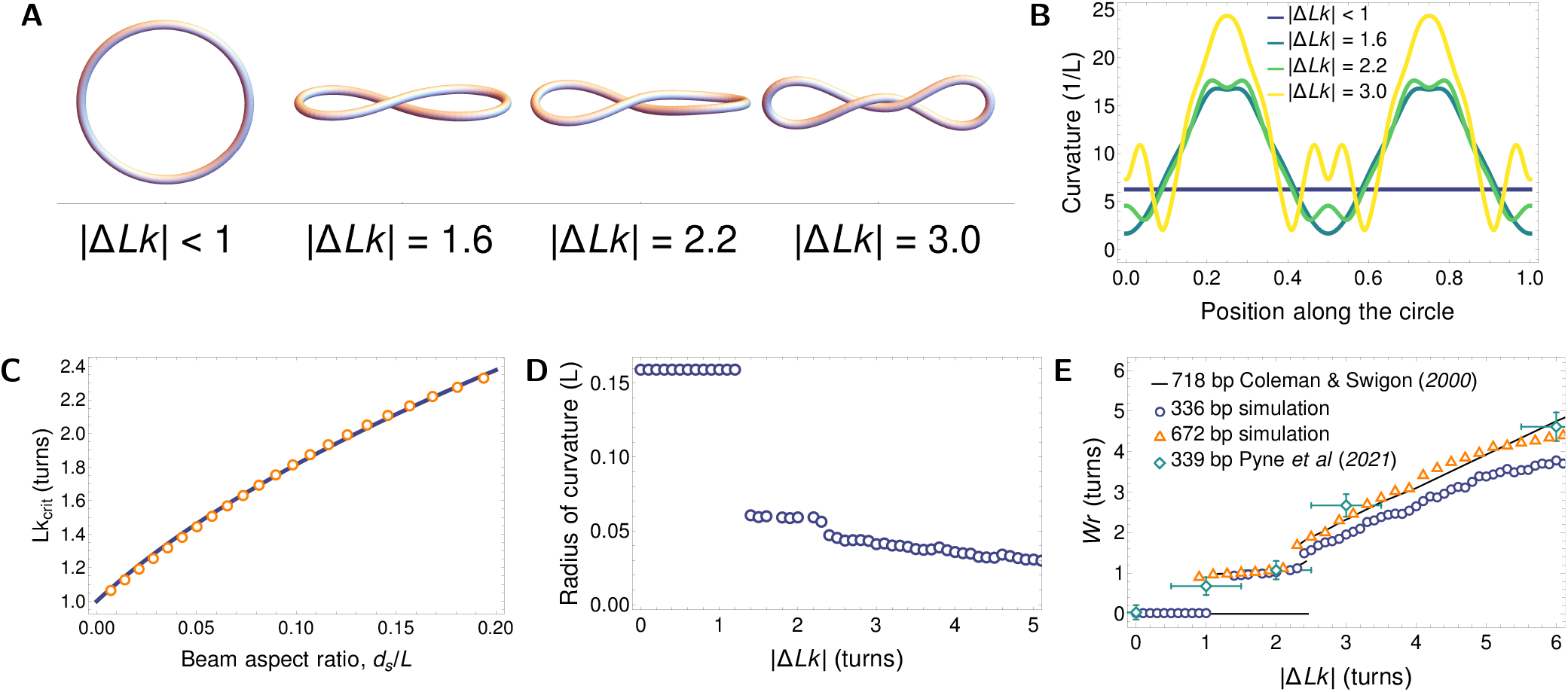
Elastic equilibrium shapes of model DNA minicircles. **(A)** Energy minimizing shapes of 336 bp minicircle of various Δ*Lk* (with *d*_*s*_*/L* = 0.018). For small |Δ*Lk*| values (< 1.6), loops adopt open circle configurations. For intermediate values (Δ*Lk* = 1.6 or 2.2), a single point of polymer contact is observed; for larger values of Δ*Lk* (> 2.2), continuous contact is observed. **(B)** Curvature distribution along the twisted loop centreline in shapes corresponding to panel **(A)**. The position along the loop is measured from the point of contact (or the centre of symmetry in multiply touching configurations). **(C)** *Lk*_crit_ above which a writhed configuration can be stable, plotted as a function of the beam aspect ratio (steric diameter to length, *d*_*s*_*/L*). Solid line shows the approximation of eqn. (**17**). **(D)** Minimal radius of curvature along the loop as a function of Δ*Lk* for the 336 bp minicircle. This radius decreases monotonically with Δ*Lk*, leading to increasing bending stresses. **(E)** Writhe of energy-minimising shapes as a function of the aspect ratios *d*_*s*_*/L* = 0.018, 0.0090, and 0.0082 for 336 bp, 672 bp, and 718 bp DNA minicircles respectively. For small values of |Δ*Lk*|, only flat (open circle) configurations are permitted but above *Lk*_crit_, one or more twists is relaxed by writhing. For large values of Δ*Lk*, around 90 % of torsional energy is relaxed by shape deformation. Our results are shown next to the continuum model predictions of Coleman & Swigon (38) and atomistic MD simulations of Pyne et al. (15).

For larger |Δ*Lk*|, increased curvature was present even at points close to the contact line. The variation of curvature along the loop centreline, shown in Figure 3C, might hint at sites of potential base pair instability or other configurational changes. The effect of localised curvature regions would be further compounded by the transmission of mechanical stress along the DNA backbone to promote DNA kinking at distant sites (13, 15, 58, 59) and base pair disruption, as reported by Fogg *et al*. (12).

The value of *Lk*_crit_ for the open circle-figure-8 transition depends on the aspect ratio *d*_*s*_*/L*. In Figure 3C, we show the dependence of *Lk*_crit_ on the aspect ratio numerically, showing that for thicker beams a transition to writhed configurations required an increased negative supercoiling. A very thin filament can be stably writhed for almost any value of |Δ*Lk*| greater than 1. However, for beams with a larger thickness, more torsional stress was required to stabilize writhed configurations and prevent them from unwrithing to a circular, open configuration.

For increased values of *d*_*s*_*/L*, we also empirically found a convenient approximate expression for

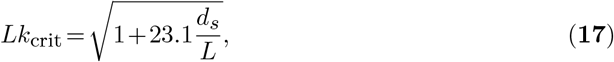

which we show in Figure 3C. To further characterize the writhed shapes, in Figure 3D we plotted the minimal radius of curvature of a twisted beam as a function of Δ*Lk*. For low values of Δ*Lk*, the constant value reflected the purely circular equilibrium. Above *Lk*_crit_, the loop became writhed, with an increased curvature. The monotonically decreasing relationship of *L/κ* showed that increasingly supercoiled beams tended to have tighter bends and, therefore, stored larger bending energy. Finally, in Figure 3E, we present the writhe of the resulting configuration, calculated with our method as a function of Δ*Lk*. The values were calculated for both lengths, 336 and 672 bp. For comparison, we also plotted the results of computations of Coleman & Swigon (38) for *L* = 718 bp as a solid line, and results of base pair-resolution MD simulations of Pyne et al. (15) for a 339 bp minicircle, confirming the observed trend. The total *Wr* as a function of Δ*Lk* seemed to be weakly dependent on the minicircle length. Writhing, therefore, emerged as a universal mechanism of stress release for twisted loops.

#### Shape and stability of supercoiled configurations

Although purely elastic considerations suggest a plethora of possible writhed configurations, we mostly observed simple minicircle conformations in the cryoET measurements of Irobalieva et al. (13), rather than more intricate shapes. This is because the latter have higher energies and are thus less frequently realized (38). For a given Δ*Lk* and at low temperatures, the shapes associated with higher energies were unfavorable compared to the ground state solution determined by our energy minimization procedure.

Equilibrium shapes measured in cryoET experiments in Ref. (13) were divided into eight groups termed, open circle, open figure-8, figure-8, racquet, handcuffs, needle, rod, and other. For a quantitative comparison, we reduced the complexity by distinguishing only between contact-free (corresponding to open minicircle configurations) and self-touching (corresponding to writhed minicircle configurations) solutions. Using the elastic beam framework, we determined the regions of stability of these solutions in terms of Δ*Lk* for a given DNA loop length. We present them in Figure 4A. The orange-shaded region close to Δ*Lk* = 0 favors an open circle as the stable configuration. For under-twisted configurations, we identified a region of multistability, where we found both open circular and figure-8 shapes. Finally, when the inherent Δ*Lk* was large enough, marked by the blue shading in the figure, the self-touching shapes became the only stable energy minimum. The stable circular shape region narrowed down from |Δ*Lk*| < 1.1 for 336 bp minicircles to |Δ*Lk*| < 1.2 for 672 bp and 718 bp minicircles.

**Figure 4.**
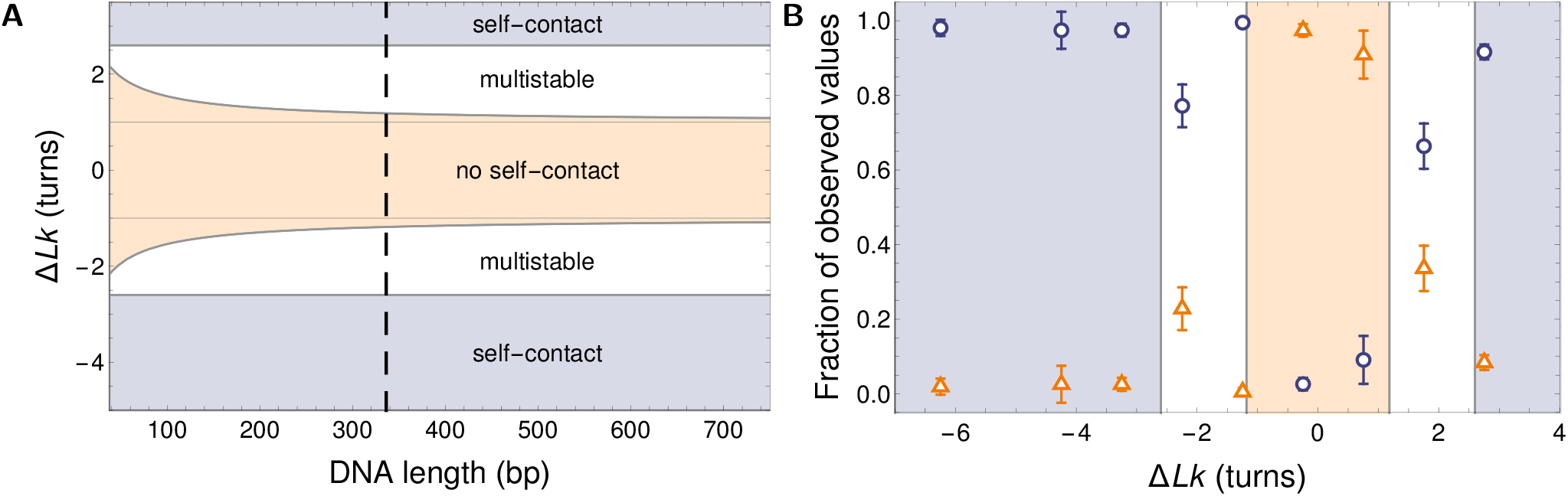
Regimes of shape stability for model DNA minicircles. **(A)** Phase diagram of different regimes of stability of twisted DNA as a function of Δ*Lk* and DNA length. The phase space is divided into three regions: only open circular configurations permissible (tan-shaded area), both circular and writhed configurations permissible (multistability; white area), and only writhed configurations permissible (blue-shaded area). The dashed line marks the 336 bp minicircles, for which experimental shape data are available. **(B)** Plot of the relative occurrence rate of configurations with (blue circles) and without (orange triangles) points of self contact, based on cryoET shapes of 336 bp minicircles measured by Irobalieva et al. (13) and overlayed on our theoretical stability predictions with the same colour coding as in panel **(A)**. Error bars represent 2 standard deviations.

For the particular length of *L* = 336 bp, in Figure 4B we compared the theoretical predictions to the cryoET data of minicircle shapes (13). We measured the fractions of open and self-touching configurations in the measured population of minicircle shapes. For Δ*Lk* close to zero, we saw that most of the shapes were open loops, with only a fraction of about 10 % showing self-contact. This situation changed with increased Δ*Lk*. In the predicted region of multistability, we saw that the fractions of writhed configurations increased, but there was still a pronounced population of open circular shapes that vanished almost completely when increasing |Δ*Lk*|. This observation is in excellent agreement with our prediction for the new stability region at |Δ*Lk*| > 2.6. In particular, the most relaxed state (closest to Δ*Lk* = 0) was slightly undertwisted (Δ*Lk <* 0). For configurations with Δ*Lk* = −0.2 and Δ*Lk* = 0.8 only open configurations were predicted while for Δ*Lk* = −2.2, −1.2 and Δ*Lk* = 1.8 we observed large conformational variability corresponding to two solutions in the uniform elasticity model. For Δ*Lk* = −4.2, −3.2, and 2.8 open configurations were no longer permitted by the uniform elasticity theory and these open configurations were largely absent in the prior cryoET measurements for these topoisomers. Interestingly, an outlier at Δ*Lk* = −1.2 was observed in the cryoET data, where we saw a surprising lack of open circular configurations, which could not be simply explained by our coarse-grained model. One potential explanation for this puzzling observation has been postulated that involves a coupling of limited base pair disruption with writhing (12).

### Predictions of hydrodynamic radius of DNA minicircles with different Δ*Lk*

#### Non-writhed configurations

Configurations with small values of Δ*Lk* adopt toroidal conformations; for those configurations, we have found a convenient expression for the hydrodynamic radius, consistent with asymptotic solutions by Johnson & Wu (42) and Johnson (60)

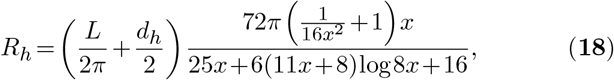

with *x* = *L/πd*_*h*_. Here, log denotes the natural logarithm. Notably, Eq. (**18**) holds sway for non-slender tori and agrees with the numerical results of Goren & O’Neill (43). For more slender tori, when *L/d*_*h*_ > 30, a simpler expression can be fitted without loss of accuracy, given by

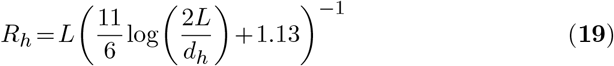

as obtained by Adamczyk et al. (61) from the numerical results of the bead-model shape approximations. Qualitatively, *R*_*h*_ is of the same order of magnitude as the experimental DNA lengths and thus it approximately scales with the mass of the DNA molecule. This scaling for approximating DNA is in contrast with globular models used for some proteins that scale with the cubic root of mass instead.

#### Writhed configurations

Configurations with intermediate values of Δ*Lk* adopt either toroidal or writhed shapes. To generate hydrodynamic predictions for DNA minicircles with larger values of Δ*Lk*, we combined elastic energy minimisation with hydrodynamic bead models. To this end, we used equilibrium shapes obtained for each Δ*Lk* within the elastic beam model described before, and produced their representation as a collection of 400 rigidly attached, overlapping spherical beads arranged such that the length and thickness of the DNA molecule were left unchanged. The bead-model was then used to calculate the hydrodynamic mobility of the conglomerate within the ZENO package (44). We estimated the error of estimation of the hydrodynamic radius to be about 5% by comparing the results of bead-model calculations to known analytical solutions for highly symmetric shapes of the model molecules.

Our modeling approach reduced the problem of finding diffusion and sedimentation coefficients to the computation of the hydrodynamic radius *R*_*h*_. Once calculated theoretically, *R*_*h*_ was used to predict the diffusion coefficient from the Stokes-Einstein relation, Eq. (**2**), provided that the viscosity of the environment *η* is known. The presented results were adjusted for the buffer viscosity. The prediction of hydrodynamic radius *R*_*h*_ for DNA of any shape required additionally the knowledge of the hydrodynamic thickness of the loop *d*_*h*_. To determine the effective thickness of the molecules (related to the existence of a hydration layer), we assumed that, as a local property, it does not depend on the shape. Then, we used the measured AUC data for nicked and relaxed (Δ*Lk* = 0) 336 bp and 672 bp minicircles, assumed to be toroidal. Knowing the dependence of the DNA hydrodynamic radius on the molecule aspect ratio from the ZENO software tool, we compared it to the AUC results for the hydrodynamic radii of the 336 bp and 672 bp DNA molecules, and we fitted the same value of *d*_*h*_ to both minicircles to reproduce their experimental hydrodynamic radii. The dependence of *R*_*h*_ on the aspect ratio is presented in Figure 5A, together with the two measured values. Fitting the theoretical curve to these two data points yielded the hydrodynamic diameter of *d*_*h*_ = 29.4 Å (corresponding to *d*_*h*_ */L* = 2.6 ×10^−2^ for the 336 bp minicircles and *d*_*h*_*/L* = 1.3 ×10^−2^ for the 672 bp minicircles). We used this value for all subsequent calculations. We note, however, that the estimation of *d*_*h*_ from diffusion measurements cannot be precise due to the logarithmic dependence of hydrodynamic parameters on this value, so the fitted value should be treated as more approximate than the number of digits provided.

**Figure 5.**
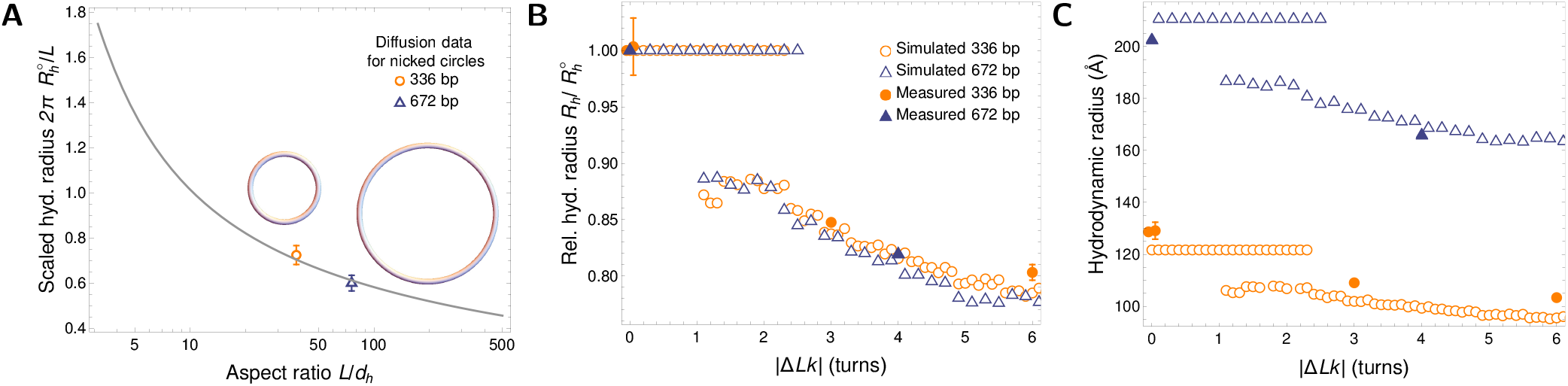
Hydrodynamic radius of DNA minicircles. **(A)** Hydrodynamic radius 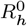 of open-circular DNA, scaled by the geometric radius *L/*2*π* of a torus, plotted for a range of DNA aspect ratios *L/d*_*h*_. Comparing the ZENO results for toroidal particles (solid line) to diffusion measurements of minicircles with 336 bp and 672 bp yields the fitted (common) hydrodynamic thickness *d*_*h*_ = 29.4 Å. We note that ZENO approximation yields high-precision results for toroidal particles (45). This value was used in all subsequent computations. Circular sketches representing molecules preserve both the relative scale and thickness. Note the logarithmic scale on the horizontal axis. **(B)** Hydrodynamic radius *R*_*h*_ of minicircle shapes, relative to the hydrodynamic radius 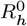 of the relaxed, open-circular shape for a range of |Δ*Lk*|. We present results of simulations (open circles and triangles) along with experimental data for 336 bp (filled diamonds) and 672 bp (filled squares) minicircles. For 336 bp (nicked); 336 bp (relaxed), and 672 bp (nicked), the simulations show a region of constant hydrodynamic radius where the shape is independent of |Δ*Lk*|. For supercoiled 336 bp containing a mixture of Δ*Lk* = −1, −2 or −3, the hydrodynamic radius is about 15 % smaller than that of an open circle. For larger values of |Δ*Lk*|, the theoretical approach seems to correctly grasp hydrodynamic radii of the resulting highly compact conformers. **(C)** Absolute values of the hydrodynamic radius *R*_*h*_ for minicircles from **(B)**, with the same point markers.

The value of excess thickness over inter-phosphate distance was significantly larger than that reported by Fernandes et al. (62) (22.8 Å)—without the details of the solvent ionic strength—However, as argued by Penkova et al. (63), the hydration shell can be as thick as 16 Å, corresponding to diameters as large as 40 Å. This value is sensitive not only to the ionic strength of the solvent, but also to the details of ion composition (64). Moreover, small deviations from the toroidal shape of the nicked and relaxed DNA shapes are expected, as caused by Brownian motion.

Figure 5B demonstrates that the difference of shapes with different Δ*Lk* is significant and similar in theory and experiments. We present therein the hydrodynamic radius for writhed configurations calculated for the two investigated DNA lengths with the same thickness (*d*_*h*_ = 29.4 Å). The plot shows the radius relative to that of a toroidal particle of the same *d*_*h*_, denoted by 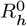. We normalize theoretical results by calculations at Δ *Lk* = 0, while experimental data are rescaled by respective results for a nicked/relaxed configuration of a minicircle of a given length. Open circles and triangles in the graph mark theoretical results. For the 336 bp minicircle, two experimental values were available, namely that of a nicked and relaxed configuration. In these cases, we rescaled the experimental data by the mean of the two radii. The values of the relative radii are equal to unity for Δ*Lk* = 0 by definition, but even the non-zero Δ*Lk* simulations predicted a region of stable circular configurations with unchanged hydrodynamic radius.

Increasing Δ*Lk* led to the emergence of highly writhed configurations, which tended to be more compact and therefore had a smaller hydrodynamic radius than the open circular configurations seen at low Δ*Lk*. We predict that topological writhing could reduce *R*_*h*_ by about 15 % in experimentally relevant conditions.

We additionally plotted available experimental results for relaxed and supercoiled minicircles, which compared favorably with theoretical predictions. The theory seems to correctly determine *R*_*h*_ obtained from AUC measurements, which opens an efficient route to calculate the hydrodynamic transport coefficients also for highly writhed conformations. In Figure 5C we also presented the same results in absolute terms. Because the fitted value of hydrodynamic thickness in Figure 5A lies between the estimates based on 336 bp and 672 bp only, we see the value of *R*_*h*_ for Δ*Lk* = 0 to be slightly overestimated by the simulation for 672 bp and underestimated for 336 bp. We emphasize that this difference is an effect of the thickness fitting procedure. This systematic difference can be caused by slightly non-toroidal DNA shapes, caused by Brownian motion.

Our AUC data, together with the theoretical modelling shown in Figure 5, yield a length-invariant observation— the ratio of *R*_*h*_ of open circular (nicked or relaxed) minicircle topoisomers to the *R*_*h*_ of compact (supercoiled) topoisomers is approximately 5:4. This value holds true for both 336 bp and 672 bp minicircles, even though the length *L* is larger than the persistence length *P* for 672 bp minicircles, with *L*≈ 4*P*. This length-invariant ratio of 5:4 holds true as long as thermal effects are negligible. This theoretical result is particularly robust because it is independent of the viscosity of the buffer or the minicircle length. We attributed this robustness to the logarithmic dependence of the hydrodynamic models on the thickness of slender bodies. Additionally, the ratio of *R*_*h*_ of 336 bp linear to *R*_*h*_ of 336 bp nicked is approximately 7:6.

These results can also be viewed in absolute, rather than relative terms (Figure 3), where the theoretical predictions for the values of the sedimentation and diffusion coefficients are displayed together with the experimental results. We obtained a good agreement for the larger of the two minicircles (672 bp) and for linear, relaxed, and nicked samples of the smaller minicircle (336 bp). For highly writhed configurations of the smaller minicircle, theoretical predictions suggest a more compact conformation than observed. This result could be attributed to the limits of applicability of the linear elasticity theory to the very tight bend radii necessary for energy minimisation in these configurations, or to a rough estimate of shapes and width of nicked and relaxed minicircles.

To test the effectiveness of our modelling approach, in Table 2 we compared the measured and predicted values of *D* and *s*. To compute *D*, data from nicked samples were used as a calibration of the only fitting parameter: the effective hydrodynamic diameter of the DNA molecule. With the use of *d*_*h*_, all the *D* values were purely theoretical predictions based on the DNA length and the value determined in separate experiments.

**Table 2.**
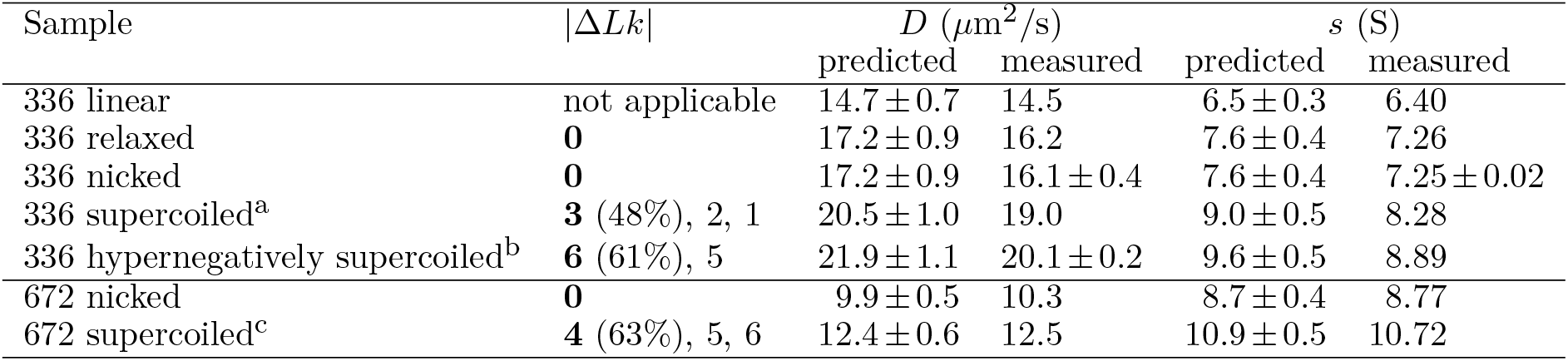
Comparison of predicted and measured diffusion and sedimentation coefficients in the buffer. ^*a*^Predictions done for the dominant species in the mix (Δ*Lk* = −3). ^*b*^Predictions done for the dominant species in the mix (Δ*Lk* = −6). ^*c*^Predictions done for the dominant species in the mix (Δ*Lk* = −4). Where experimental errors are not shown, the confidence limits from the global Monte Carlo analysis of the AUC data were exactly 0. We roughly estimated the errors of theoretical values as 5 %. Sedimentation coefficients are measured in svedbergs, with 1 S= 10^−13^ s. Experimental confidence limits are reported only when the last digit of the result would be affected.

Because supercoiled and hyper-supercoiled samples contain mixtures of minicircles with different Δ*Lk*, theoretical predictions are given for the most common species in a mixture. These configurations are sketched in Figure 6. To calculate *s*, we used the approximately constant value of the PSV determined from the AUC experiments. In all cases, the deviation between theory and experiment was below 8 %. The simplified hydroelastic model provides practical estimates of the experimentally accessible quantities, and thus may be used to discern populations of minicircles differing in Δ*Lk*.

**Figure 6.**
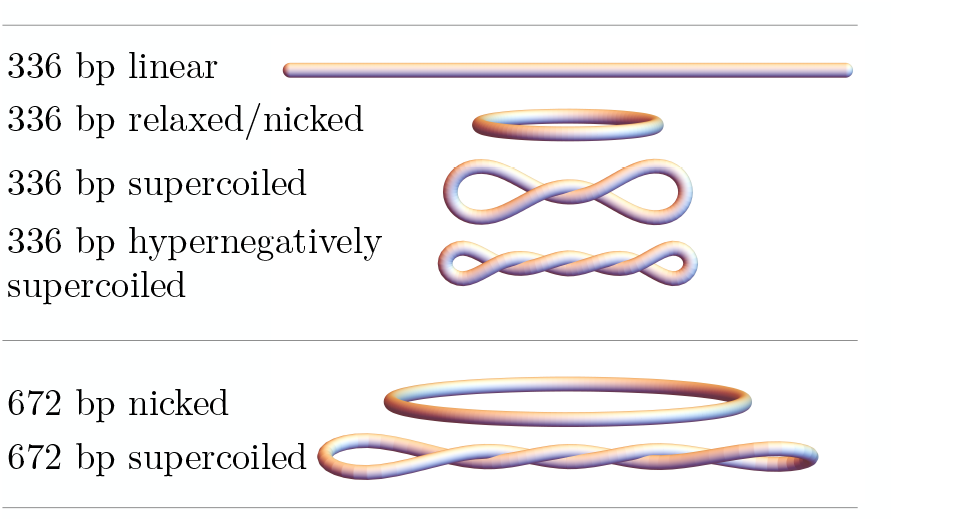
Sketches of model shapes in a given minicircle configuration used for hydrodynamic simulations with Δ*Lk* specified in the caption of Table 2. The sketches have realistic aspect ratios (*d*_*h*_*/L*) and preserve the relative size.

Our modelling experiments reveal that the change of Δ*Lk* by one turn (for example from Δ*Lk* = −4 to Δ*Lk* = −5) changes the frictional ratio by ∼2 %— a value similar to theory-experiment deviations and similar to the noise level coming from the Monte Carlo optimizations of energy minimizing shapes. Such small gradients could be the reason why AUC measurements cannot distinguish topoisomers which differ in Δ*Lk* by only a single turn, especially if the concentrations of the different topoisomers in the mix differ significantly. The samples analyzed by AUC contained a mixture of topoisomers that behaved as a single species because the major species in each sample were within one Δ*Lk* of each other.

## DISCUSSION

Methods that measure the hydrodynamic radius *R*_*h*_, such as gel electrophoresis and diffusion-sedimentation AUC experiments can be qualitatively described as sorting molecules by their size. Unlike for gel electrophoresis, however, when assessing diffusion via AUC, size contributions can be separated from viscosity, temperature, and PSV by the appropriate scaling, which yields *R*_*h*_.

In electrophoresis, molecules squeeze their way through pores in the gel matrix. One of the major differences between agarose and polyacrylamide is the average pore size, with agarose gels typically having larger sized pores (65). In typical electrophoresis experiments, the primary determinant of the migration speed is the molecular weight and charge of the molecule. This is in contrast with our observations. Here, the DNA molecule of a given length has a fixed charge and molecular weight, but the speed of migration through the matrix of small pores like the polyacrylamide gel varies greatly when shape of the molecule is changed. Comparing gel results to the AUC data allows us to confidently say that knowing the molecule’s *R*_*h*_ is insufficient to predict its electrophoretic mobility as the gel electrophoresis and AUC had separated the minicircle topoisomers differently. Supercoiling influences minicircle electrophoretic mobility much more than hydrodynamic mobility (as measured by AUC); the same change in supercoiling increases electrophoretic mobility by 400 % while increasing mobility only by 20 %. Differences between the methods can be even more dramatic, for the same experimental conditions, the linearized 336 bp minicircle showed 600 % increase of electrophoretic mobility as compared to the nicked or relaxed form, at the same time having 14% *smaller* hydrodynamic mobility compared to nicked or relaxed as measured by AUC.

Introducing circularization presents a significant obstacle to extending quantitative electrophoresis methods, such as one proposed by Ziraldo et al. (66). Calibrating gel electrophoresis measurements with a ladder of relaxed or nicked circular DNA of different lengths would be insufficient, since the contribution of the degree of supercoiling strongly affects the apparent DNA mass derived from the method of Ziraldo et al. (at least by a factor of two). These difficulties are further compounded by the dependence on the absolute value of applied electrophoretic-mediated force, as reported by Iubini et al. (67). For large electrophoretic-mediated forces, linear DNA is expected to migrate faster than the circular form, which is opposite to what occurs via AUC, while for small forces, the circular form migrates faster than the linear, which is the same as in AUC even when the gel properties are kept constant. One possible explanation for the differences between electrophoresis and AUC is that as the DNA is pulled through the pores of the gel matrix, it has to change its shape (this would be most significant for the Δ*Lk* = 0 samples) whereas there is no sieving in AUC. AUC, therefore, better reflects the solution properties of DNA, while gel electrophoresis offers higher resolution for separation.

Diffusion-sedimentation measurements provided by AUC are free of these obstacles and allow accurate hydrodynamic modelling. To elucidate the shapes and properties of DNA minicircles, we proposed a coarse-grained model that represents the DNA as an elongated, uniform elastic beam. Their predefined linking numbers corresponded to the degree of supercoiling. By minimising the elastic energy of a beam with a given superhelical density, we were able to find equilibrium shapes of the minicircles. We note that our coarse-grained models of minicircles are oblivious to their sequence and do not exploit information on sequence-dependent elastic properties. However, the uniform beam model of DNA elasticity presented here predicts shapes of DNA minicircles (shown in Figures 3A and 6) that compare favourably with the direct observations of 336 bp minicircles published earlier by Irobalieva et al. (13) (shown in Figure 4B). Our theoretical model predicts a very weak dependence of the shapes of DNA minicircles with a few hundred base pairs on the aspect ratio *d*_*s*_*/L*, as shown in Figure 4A. According to the model, the change in Δ*Lk*, alone, gives sufficient information to describe basic features of the minicircle configuration: Δ*Lk* = 0 is an open circle; Δ*Lk* = *±*1 is at the transition between open circle and writhed conformations; Δ*Lk* = *±*2 is multi-stable; and |Δ*Lk*| > 3 always exhibits self-contact (compare with the corresponding shapes in Figures 3A and 6). This means that no matter whether the minicircle was 336 or 672 bp, the loss of three helical turns was enough to disallow the open circle conformation as an energy-minimal solution. We conclude that it is the absolute value of Δ*Lk* and not the superhelical density *σ* =Δ*Lk/Lk*_0_ that governs the conformational landscape of small DNA minicircles (with the length having a small influence via the *d*_*s*_*/L* ratio). This finding provides an important input to future models of circularised polymers. Incorporating torsional interactions is more difficult than just bending interactions and is often neglected when constructing models of circularized molecules (68, 69, 70, 71). Since the torsional forces play a role even when Δ*Lk* is close to zero, additional care needs to be taken when generalising such models for the context of supercoiled DNA. We have shown that regardless of the length of the DNA molecule the torsional forces are of the same order of magnitude as pure bending forces.

The elastic-energy minimising shapes found for minicircles with different Δ*Lk* can have different thickness; its value was determined by calibration based on AUC experimental data. These shapes were then used to construct hydrodynamic models to compute *R*_*h*_, which then in turn was used to compare the theoretical diffusion and sedimentation coefficients with AUC measurement. The comparison (Table 2 shows general agreement which is satisfactory given the simplicity of the underlying coarse-grained model. These results confirm the predictive capabilities of uniform elasticity models, combined with a hydrodynamic calculations package (ZENO) to interpret and guide AUC measurements. We expect that the proposed modelling strategy could be beneficially employed to similar problems in the dynamics of DNA and perhaps extended further to account for sequence-specific effects and modified intramolecular interactions.

This work represents significant progress in understanding and modelling the sedimentation of a biological molecule with a complex and dynamic conformation. Sedimentation of roughly spherical molecules (e.g., many proteins) is fairly well understood, but neither linear nor supercoiled DNA adopts a spherical conformation. Using 336 bp minicircles as a model system we are now able, for the first time, to fully test the theoretical models. We anticipate that this will allow us to further improve these models and expand the use of AUC to include a larger repertoire of important and complex biological molecules. We have successfully applied two stage modelling (combination of energy minimization to find the shape then applying hydrodynamic modelling for rigid configurations) to find hydrodynamic properties of a DNA molecule. In the context of future work, presented approximate calculations of relative importance of different physical phenomena on the energetic landscape governing the molecules shape can be easily replicated and the efficacy of two stage modelling can be quickly assessed beforehand. Similarly the limits of applicability of the current approach are determined by the persistence length, the Debye length, and the molecule length alone.

## CONCLUSION

The work presented here is a step toward understanding how supercoiling and curvature affect DNA shape and hydrodynamic properties, which in turn affect important DNA activities. The DNA solvation shell and counterions as well as DNA shape all affect DNA structure to influence how the DNA code is protected, accessed, modified, and activated. It will take a combination of approaches to fully understand this remarkable molecule.

## DATA AVAILABILITY

The UltraScan software used to analyze the AUC data is open source and freely available from [Github repository](https://github.com/ehb54), the AUC data itself is available upon request from the [UltraScan LIMS server](https://uslims.uleth.ca/uslims3_CCH/) at the Canadian Center for Hydrodynamics. Algorithm for finding minimal energy shapes, its initial conditions and final configurations can be found in Github repository RadostW/twisted-loop. *Lk* −*Wr* pairs computed from energy minimal shapes as well as hydrodynamic radii computed using the ZENO software can be found in Github repository RadostW/twisted-dna-results.

## ACKNOWLEDGEMENTS

The Authors thank Prof. De Witt Sumners and Dr. David Swigon for helpful remarks. The work of RW and ML was supported by the National Science Centre of Poland grant Sonata to ML no. 2018/31/D/ST3/02408. This work was also supported by the Canada 150 Research Chairs program (C150-2017-00015, BD), the National Institutes of Health (1R01GM120600, BD) and the Canadian Natural Science and Engineering Research Council (DG-RGPIN-2019-05637, BD). The Canadian Center for Hydrodynamics is funded by the Canada Foundation for Innovation (CFI-37589, BD). UltraScan supercomputer calculations were supported through NSF/XSEDE grant TG-MCB070039N (BD), and University of Texas grant TG457201 (BD). MLEJ was supported in part by National Science Centre of Poland under grant UMO-2018/31/B/ST8/03640. The authors acknowledge funding (to LZ) from National Institutes of Health grant R35 GM141793 and National Science Foundation grant MCB 2107527.

---

**Supplementary Table 1.** Topological composition of the DNA samples used. The composition of the samples was determined by quantification of fluorescently stained gels using image analysis software. N.D.: not detected.

| Minicircle length, bp | Sample | Topoisomers present (% of total) | Trace species (% of total) |
| --- | --- | --- | --- |
| 336 | “Supercoiled” | $\Delta Lk = -3$ (48%)<br>$\Delta Lk = -2$ (41%) | $\Delta Lk = -1$ (7 %)<br>Nicked 336 bp (1 %)<br>Supercoiled 672 bp (3 %) |
|  | Nicked | Nicked (93 %) | Nicked 672 bp (7 %) |
|  | Relaxed | Relaxed (95 %) | Relaxed 672 bp (5 %) |
| | “Hypernegatively supercoiled” | $\Delta Lk = -6$ (61 %)<br>$\Delta Lk = -5$ (33 %) | Nicked 336 bp (4 %)<br>Supercoiled 672 bp (3 %) |
|  | Linear | Linear (100 %) | N.D. |
| 672 | “Supercoiled” | $\Delta Lk = -4$ (63 %)<br>$\Delta Lk = -5$ (24 %) | $\Delta Lk = -6$ (6 %)<br>$\Delta Lk = -2$ (4 %)<br>$\Delta Lk = -3$ (2 %)<br>Nicked (2 %) |
|  | Nicked | Nicked (100 %) | N.D. |

**Supplementary Figure 1.**
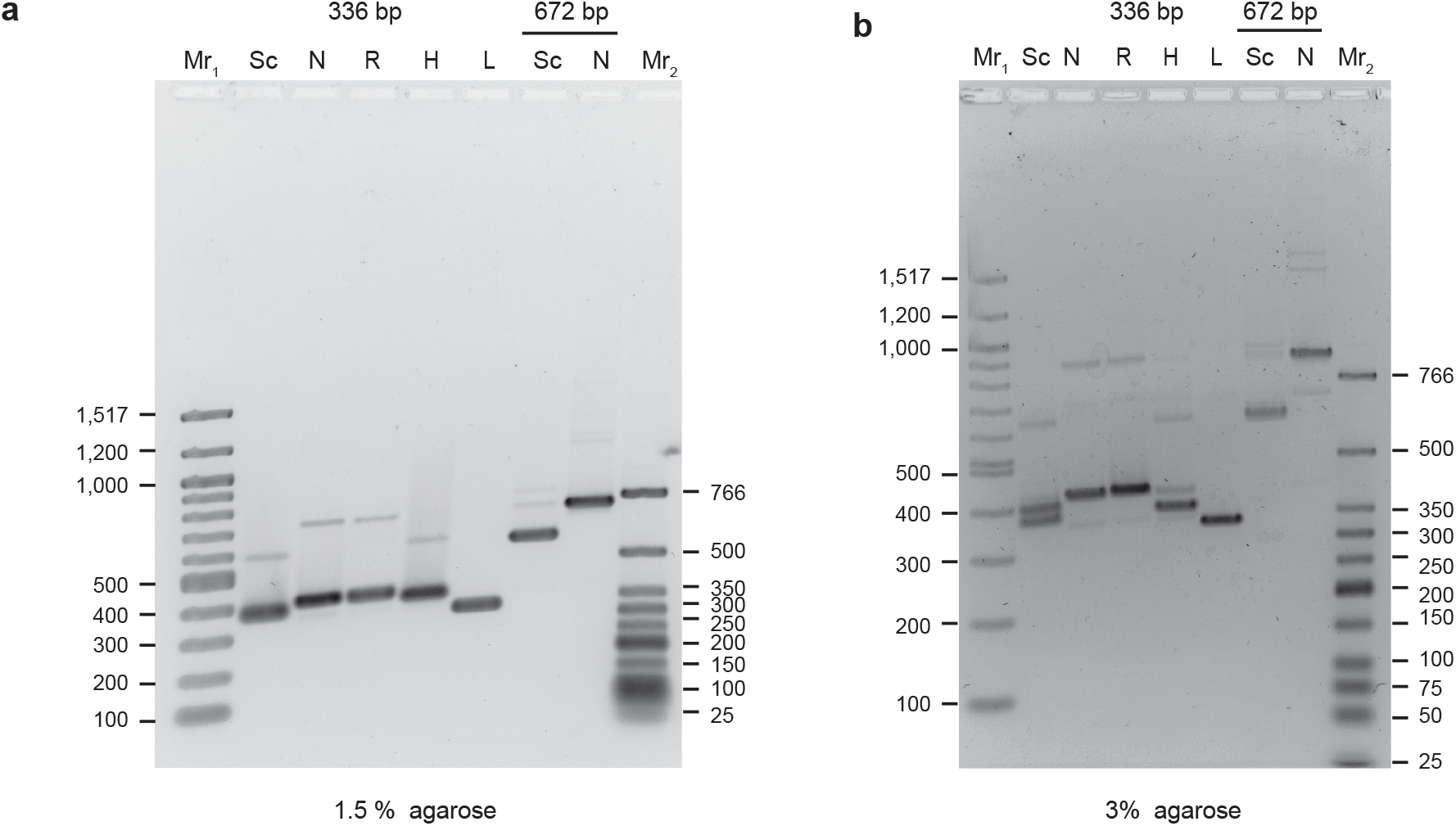
Electrophoretic mobility of minicircle DNA on agarose gels. DNA samples were analyzed by agarose gel electrophoresis in either a) 1.5 % agarose or b) 3 % agarose in the presence of 1 mM EDTA. Mr_1_: 100 bp DNA ladder, lanes 2–6: 336 bp minicircle DNA samples (Sc: “supercoiled”, N: nicked, R: relaxed, H: “hypernegatively supercoiled”, L: linear), lanes 7–8: 672 bp DNA samples (Sc: “supercoiled”, N: nicked), Mr_1_: low molecular weight DNA ladder.

